# Unravelling the developmental roadmap towards human brown adipose tissue

**DOI:** 10.1101/2020.07.09.194886

**Authors:** Stefania Carobbio, Anne-Claire Guenantin, Myriam Bahri, Isabella Samuelson, Floris Honig, Sonia Rodriguez-Fdez, Kathleen Long, Ioannis Kamzolas, Sherine Awad, Dunja Lukovic, Slaven Erceg, Andrew Bassett, Sasha Mendjan, Ludovic Vallier, Barry S. Rosen, Davide Chiarugi, Antonio Vidal-Puig

**Author notes:** Corresponding authors: Stefania Carobbio, and Antonio Vidal-Puig. These authors contributed equally to the work (Stefania Carobbio, Anne-Claire Guenantin).

## Abstract

Increasing brown adipose tissue (BAT) mass and activation has been proposed as a potential therapeutic strategy to treat obesity and associated cardiometabolic complications. Given that obese and diabetic patients possess low amounts of BAT, an efficient way to expand their BAT mass would be necessary if BAT is to be useful. Currently, there is limited knowledge about how human BAT develops, differentiates, and is optimally activated. Moreover, to have access to human BAT is challenging, given its low volume and being anatomically dispersed. These constrain makes detailed mechanistic studies related to BAT development and function in humans virtually impossible. To overcome these limitations, we have developed a human-relevant new protocol for the differentiation of human pluripotent stem cells (hPSCs) into brown adipocytes (BAs). Unique to this protocol is that it is chemically-defined to recapitulate a physiological step-by-step developmental path of BAT that captures transient paraxial mesoderm and BAT progenitor states, on its way to reaching the adipocyte stage finally. These hPSC-derived BAs express brown adipocyte and thermogenic markers, are insulin sensitive, and respond to β-adrenergic stimuli. This new protocol is a scalable tool to study human BAs during development.

## Introduction

Obesity and its associated cardiometabolic complications are important for global public health problems. Despite decades of intense research elucidating the mechanisms controlling energy balance and body weight, it is demoralising that probably the most effective therapy to counteract obesity is still bariatric surgery – This approach is invasive, irreversible and not devoid of risk. Given the importance of the obesity epidemic, its relevance as a risk factor for cardiometabolic complications, the severity of infectious diseases, the associated human suffering and economic burden to society, there is an urgent need for alternative safe, efficient and cost-effective solutions to combat weight gain and its related comorbidities.

Brown adipose tissue (BAT) is a thermogenic organ able to dissipate energy in the form of heat through regulated mitochondrial uncoupling. BAT thermogenic activity requires activation of a signalling cascade typically initiated by β-adrenergic stimulation. The critical effector of thermogenesis is the mitochondrial uncoupling protein 1 (UCP1), whose function is to allow protons to short-circuit the inner mitochondrial membrane, bypassing ATP synthase and dissipating the proton motive force as heat ^1^. In rodents and other small mammals, the principal role of BAT is to maintain body temperature. Moreover, sustained activation of BAT frequently leads to weight loss, presumably by promoting energy dissipation resulting in negative energy balance ^2,3^. From a homeostatic point of view, increasing BAT function would be expected to increase food intake to match the level of energy expenditure. However, there appears to be an unidentified mechanism by which the increase in food intake is attenuated ^4^. This dissociation between food intake and energy expenditure regulation provides a therapeutic window leading to net weight-loss, which, if exploited in humans, could lead to a desirable clinical outcome.

BAT is most readily found in small mammals and newborn humans ^4^, both having in common their high surface/volume ratio, which requires increased heat production for thermal homeostasis beyond what may contribute the byproduct of other metabolic processes. Imaging studies conducted in adult humans have demonstrated that BAT is present and functional in most young, lean human adults, particularly when exposed to cold ^5^. Unfortunately, obese and diabetic humans appear to have less BAT ^6,7^. The lack of BAT in these patients is, at least partially, reversible by exposure to lower temperatures ^8^. The re-appearance of BAT under this environmental condition indicates that these patients should have BAT precursors in their adipose tissue and that they could still benefit from strategies promoting the differentiation of these precursors into mature adipocytes that could subsequently be activated. This focus on the differentiation of precursors is the fundamental reason to elucidating the different stages of human BAT development. As demonstrated in rodent models, where increasing BAT mass and activation improve diabetes and dyslipidaemia ^8^, being able to increase BAT mass and activation is an attractive and, most importantly, safe therapeutic strategy.

Before BAT activation or differentiation can be considered a feasible target for therapeutic strategies, there are two significant hurdles to be overcome. The first is to understand the mechanisms controlling the mass of human BAT. While detailed information available from murine model systems provides essential insights, there are already known significant differences between rodent and human BAT regarding thermogenic capacity, marker gene expression, and importantly, dissimilar pharmacological responsiveness ^9-11^. Furthermore, the exact developmental origin and cell fate decisions determining canonical BAT development in humans are unclear. This lack of fundamental knowledge is in part due to the second constraint, the difficulty in obtaining a sufficient quantity of enough quality of human BAT. Getting human BAT requires invasive surgery, and while there are a few well characterised human BA cellular models, they are immortalised cells, and the degree to which they mimic the *in vivo* situation is unknown ^12-14^.

In addition to the issues surrounding human BAT cell lines, the available cellular and developmental models of human brown adipogenesis are certainly suboptimal ^12-14^ by their limited accessibility and severe constraints on how long they can be passaged. These impose significant limits on scalability. Overcoming these deterrents, *in vitro* differentiation of hPSCs is a promising model for the study of human BAT development, brown adipogenesis, and mature BA function. Current protocols of hPSC differentiation into BAs fall into two categories. The first type relies on derivation of either mesenchymal stem cells (MSCs) or embryoid bodies ^15-17^ before applying a chemical adipogenic stimulus ^16,17^. The second type relies on ectopic overexpression of genes that drive the BA programme ^18,19^. Both types of protocols bypass key intermediate pathways, making them unsuitable for elucidating the developmental and adipogenic as well as thermogenic signalling events physiologically leading to BAT formation. In our opinion, a gap of knowledge that is essential to address if BAT is going to be of any therapeutic use.

Here we report the development of a robust chemically defined protocol for the differentiation of hPSCs into BAs that is upscalable. Our method recapitulates the physiological roadmap to make human BAT development, by directing the pluripotent stem cell state towards the paraxial mesoderm and BA progenitor stage before finally going through adipogenic and functional maturation. This cellular model represents a unique tool to dissect the molecular mechanisms regulating human BAT development and progenitor differentiation.

## Results

### A chemically-defined protocol for differentiation of hPSCs into paraxial mesodermal progenitors

BAs and skeletal muscles arise from paraxial mesoderm, indicating a common origin for these two cell lineages ^20^. Thus, we developed a differentiation protocol that first directed pluripotent stem cells toward a mesoderm identity, then to a BAT progenitor state and finally into mature BAs (Graphical abstract and Fig.S1A). To confirm the specificity and reproducibility of this protocol, we validated it in two independent cell lines, the human embryonic stem cell line (hESC), H9 and the human induced pluripotent stem cell line (hiPSCs) KOLF2-C1 (Fig.S2). The similarity in the transcriptome of the two cell lines at each stage was confirmed by Principal Component Analysis (PCA) and clustering analysis. The remarkable similarity in both, the PCA and heatmap clustering of the two cell lines, given their differences in origin and genetic background, was purely based on the stage of development rather than the intrinsic differences between the cell lines (Fig.S1B-C). To obtain early mesodermal progenitors from undifferentiated PSCs, we cultured hPSCs for 48h (day (D)0 to D2) in Chemically Defined Medium (CDM) supplemented with insulin, FGF2 and Chiron (GSK3 inhibitor). As expected, at D2 we observed the transient upregulation of the mesodermal marker *BRACHYURY* (*TBOX*) as assessed by qPCR, immunoblot, and immunocytochemistry (Fig.1A-C and Fig.S3A-B). The high proportion of TBOX positive cells indicates the optimised efficiency of the protocol of differentiation to mesodermal precursors (Fig. 1C and Fig.S3B). Over the same period, expression of pluripotency markers such as *NANOG, SOX2*, and *OCT3/4*, all decreased (Fig.S2A). Gene Set Enrichment Analysis (GSEA) of the RNA-seq data comparing D4 to D0 confirmed the generation of mesoderm-like progenitors (Fig.1D and Fig.S3C).

**Figure 1.**
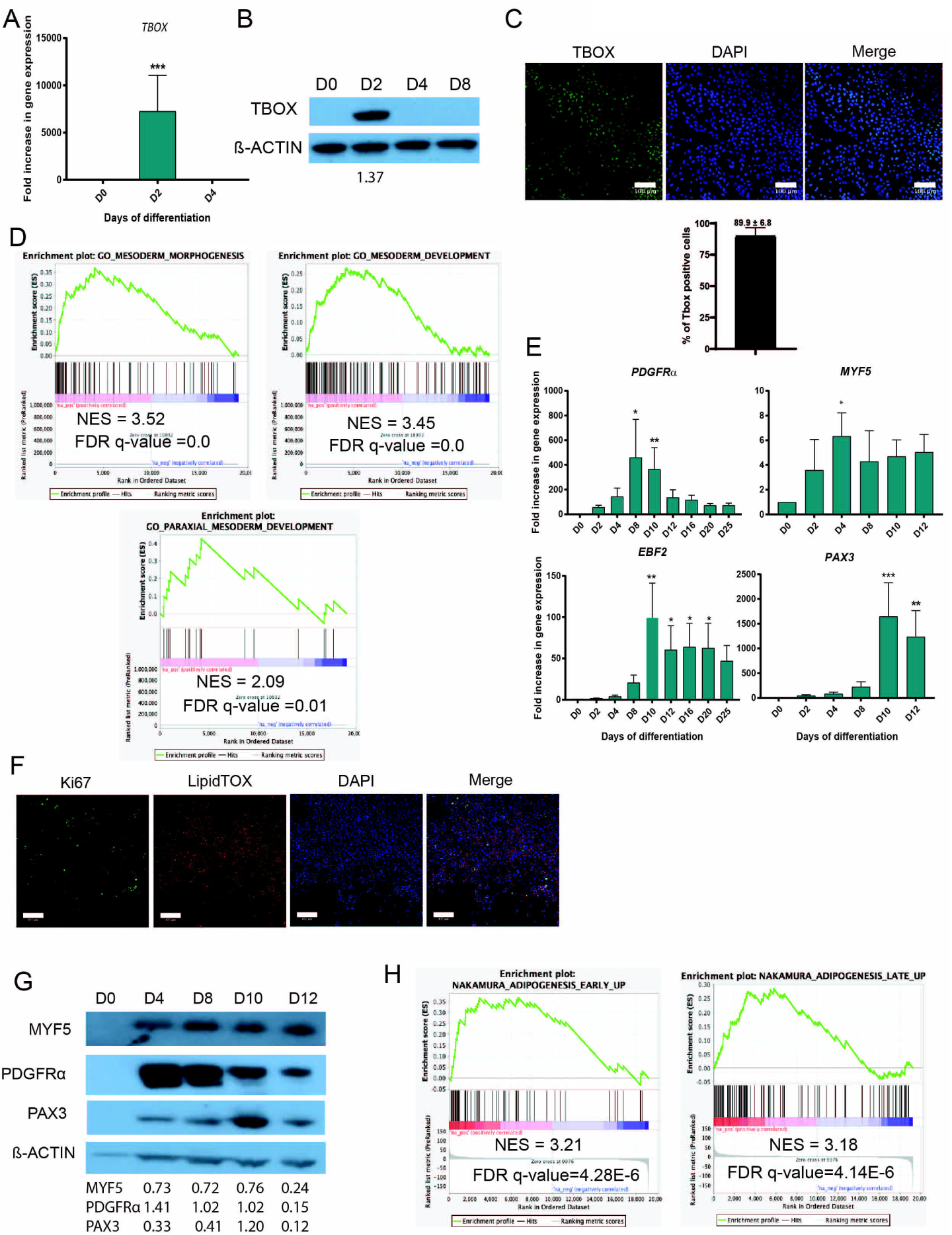
Differentiation of induced pluripotent stem cells into mesoderm and adipose progenitors. (A) RT-qPCR analysis of expression of the mesodermal marker *TBOX* (mean ± SEM arbitrary units (A.U.) relative to D0; n ≥ 3 independent experiments; ***p<0.001 relative to D0). *GAPDH* was used as the housekeeping gene. (B) Detection of TBOX from D0 to D8 by western blot (WB). β-ACTIN was used as loading control. Western blot quantification is shown underneath the WB image. (C) TBOX immunostaining of mesodermal progenitors at D2 (Green). Nuclei were stained with DAPI (blue). Bars: 100 µm. TBOX positive cells were quantified using CellProfiler (SEM ± mean, n=3 technical replicates). (D) Gene Set Enrichment Analysis of H9-derived cells on D4 *vs*. D0 using GSEA (n=3 independent experiments), using the “mesoderm morphogenesis” GO:48332, “mesoderm development” GO:0007498 and “paraxial mesoderm development” GO: 0048339 datasets. (E) RT-qPCR analysis of expression of indicated genes from D0 to D12 (mean ± SEM A.U. relative to D0; n ≥ 3 experiments; *p<0.05, **p<0.01 and ***p<0,005 relative to D0). *GAPDH* was used as housekeeping gene. (F) Ki67 immunostaining of adipose progenitors at D12 (green). Nuclei were stained with DAPI (blue). LipidTOX was used to stain the lipid droplets (red). Bars: 100 µm. (G) Detection of MYF5, PDGFRα, and PAX3 in H9 on D0, D4, D8, D10, and D12 by western blot. β-ACTIN was used as loading control. Western blot quantification is shown underneath the WB image. (H) Gene Set Enrichment Analysis of H9-derived adipose progenitors cells using published datasets (“Nakamura adipogenesis early up” and “Nakamura adipogenesis late up”), with early and late adipogenesis transcriptomic signatures on D12 *vs* D0, compared to human adult adipose stromal cell signature (n = 3 independent experiments).

Paraxial mesoderm arises from the primitive streak ^21^ and gives rise to different cell layers, including the dermomyotome ^22^ from which BAT and skeletal muscle ^23^ derive. From D2 to D4 we incubated the cells in CDM and added insulin, FGF2, and retinoic acid (RA) to drive the cells to a paraxial mesoderm-like stage. The combination of some of these factors has been reported to induce paraxial mesodermal differentiation from hPSCs *in vitro* ^24^. Using this induction cocktail, we obtained progenitor cells expressing mRNA and protein of the typical paraxial mesoderm markers *MYF5* and *PDGFRα* 25 (Fig.1E, 1G and Fig.S3A, S3D). Moreover, GSEA analysis of the RNA-seq data generated from these differentiating cells at D4 compared to D0 confirmed the generation of paraxial mesoderm-like progenitors *in vitro* (Fig.1D). These results confirmed that our protocol promoted the development of paraxial mesodermal precursors from undifferentiated hPSCs ^21^.

### Differentiation of paraxial mesodermal precursors into brown adipocyte progenitors

At D4, paraxial mesoderm-like precursors became brown adipocyte precursor cells upon treatment for 48h with insulin, FGF2, Chiron, and LDN-193189 (a BMP inhibitor). Also, ascorbic acid was used to promote proliferation ^26^. Following adipocyte precursor formation, the cells were treated with adipogenic induction between D6-D8, requiring culturing cells in DMEM/HAMF12 medium supplemented with triiodothyronine (T3), dexamethasone, 3-isobutyl-1-methylxanthine (IBMX), biotin, pantothenate, insulin, rosiglitazone, ascorbic acid and serum. Under these conditions, the progenitors proliferated, as confirmed at D12 by immunocytochemistry with proliferation marker Ki67 (Fig.1F and Fig.S3F). Following four days of adipogenic induction (D12), cells expressed the BA lineage markers *PAX3* and *EBF2* (Fig.1E, 1G, and Fig.S3A). GSEA analysis of RNA-sequencing data comparing D12 and D0 showed a high degree (with a Network Enrichment Score (NES) > 3.1) of similarity in the gene expression signature of our cells and primary human stromovascular cells isolated from adipose tissue 27 (Fig. 1H and Fig.S3E).

Thus, these results indicate that adipogenic induction of hPSC-derived mesodermal precursors drives the emergence of proliferating adipocyte progenitors.

### Generation of human adipocytes expressing classical adipose markers

After adipogenic induction, the PSC-derived brown precursors were cultured in the presence of T3, dexamethasone, biotin, pantothenate, insulin, rosiglitazone and oleate from D12 onwards. At this stage, these cells exhibited increased mRNA expression of adipogenic markers such as *C/EBP*α, *C/EBPβ, C/EBPδ* and *PPARγ* (Fig.2A), lipid droplet proteins including *ADRP1* and *PLIN1*, as well as the fatty acid transporter *CD36* ^28^(Fig.3A). Of note, *C/EBP*β expression was not transitory, as in white adipogenesis, but sustained, suggesting a more “brown” adipogenic program. Immunocytochemical detection of C/EBPα, ADRP1 and PLIN1 was observed in cells positive for LipidTOX (Fig.2B, 3Band Fig.S3F, S4A), suggesting that cells showing lipid accumulation were undergoing *bona fide* adipogenesis. The adipose tissue progenitors positive for C/EBPα represented 51.3% of the cells (Fig.2B). In line with this, LipidTOX intensity levels, mean lipid area per cell, and the number of LipidTOX positive cells gradually increased from D0 to D25 of differentiation (Fig. 3C-D and Fig. S3G). As expected, the proportion of lipid-containing cells GSEA analysis of our PSC-derived adipocytes at D25 showed a strong correlation with the Gene Ontology (GO) pathways related to “fat differentiation", “lipid homeostasis” “lipid metabolic processes” and “lipid catabolic processes”, indicating the development of cells with a coordinated lipid metabolic program (Fig.2C and Fig.S4G). Comparable results were observed performing a similar analysis with the PAZ6 human BAs transcriptome at D14 (mature adipocytes) *vs* D0 (preadipocytes), confirming the similarity of hPSC-derived BAs to human BAs (Fig.S5).

**Figure 2.**
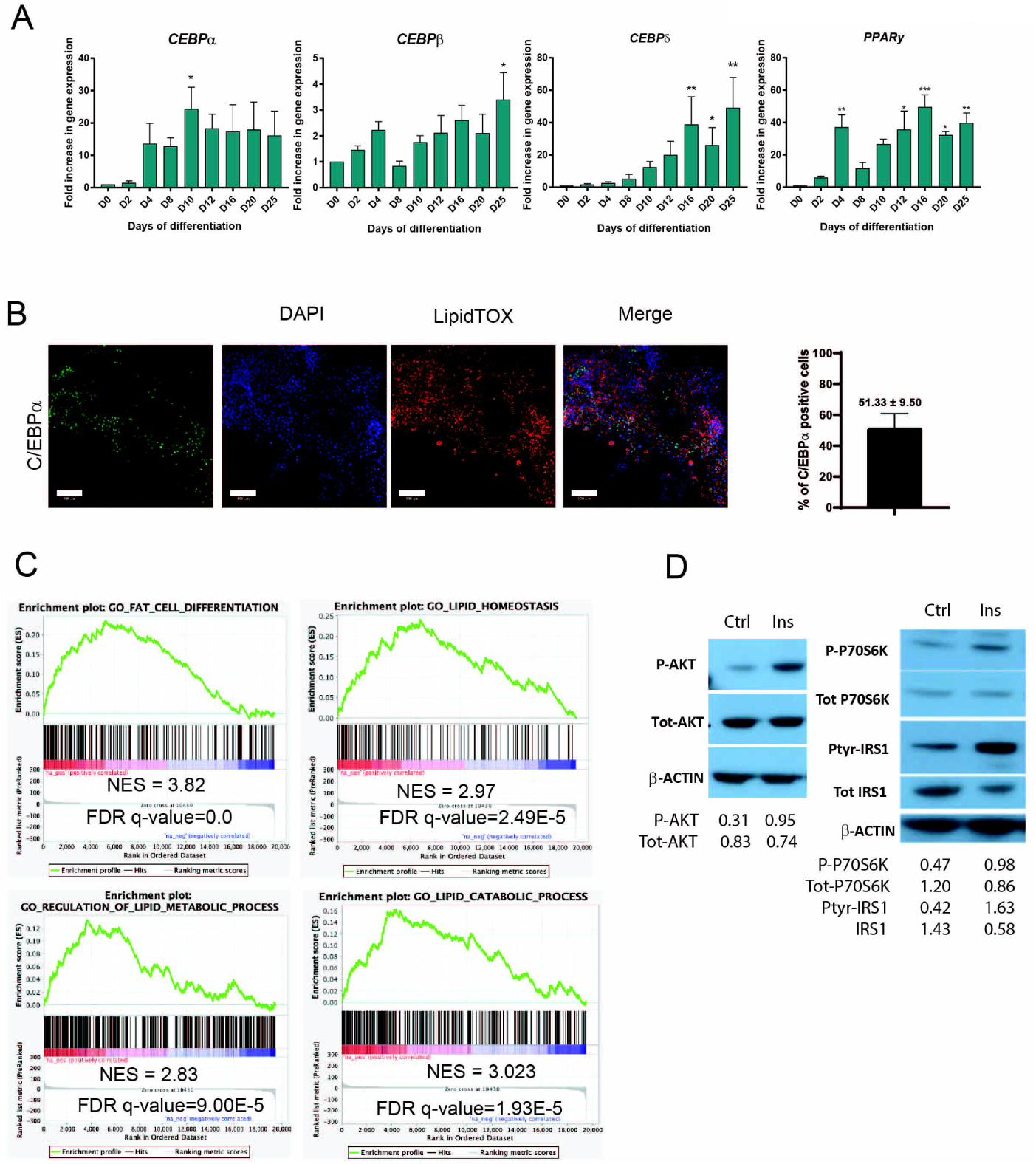
Brown adipocytes progenitors differentiation into adipocytes. (A) Time-course mRNA expression of indicated adipocytes transcription factors during differentiation of H9 (n ≥ 3 experiments; mean ± SEM A.U. *p<0.05, **p<0.01 and ***p<0,005 relative to D0). GAPDH was used as housekeeping gene. (B) Immunodetection of C/EBPα (green) in lipid-containing adipocytes (LipidTOX - red) (H9) on D20. Nuclei were stained with DAPI. Bars: 100 µm. C/EBPα positive cells were quantified using CellProfiler (SEM ± mean, n=3 technical replicates). (C) Gene Set Enrichment Analysis of H9-derived adipose cells at D25 *vs* D0 with GO datasets (“fat cell differentiation” GO:0045444, “lipid homeostasis” GO:0055088, “regulation of lipid metabolic process” GO:0006629 and “lipid catabolic process” GO:0016042), (n = 3 independent experiments). (D) Short-term insulin-mediated phosphorylation of AKT, IRS1 and P70S6K in H9 on D25. β-ACTIN was used as loading control. Western blot quatification is shown underneath the WB image.

**Figure 3.**
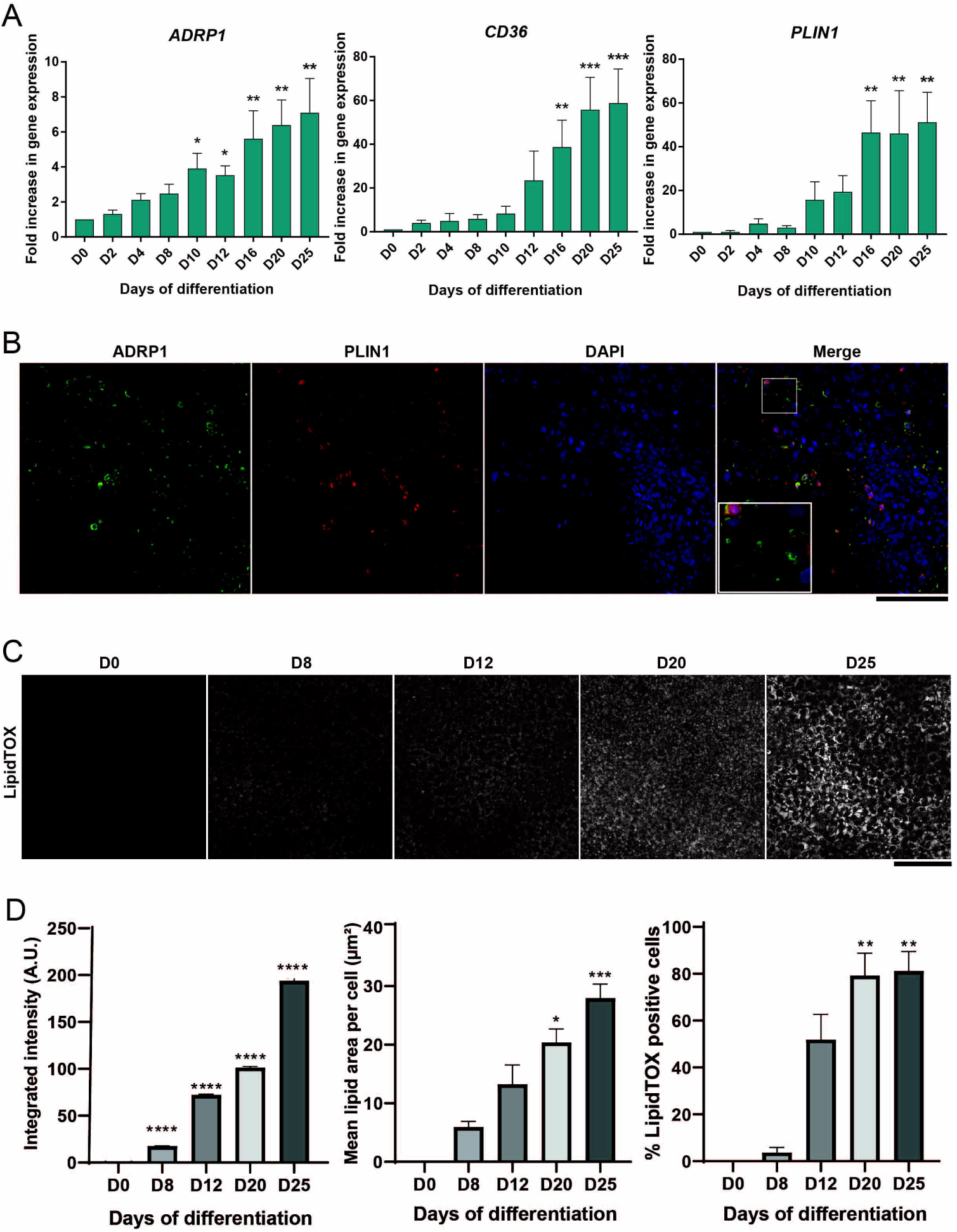
Brown adipocytes lipid accumulation. (A) Time-course mRNA expression of indicated adipocytes key factors involved in lipid accumulation during differentiation of H9 (n ≥ 3 experiments; mean ± SEM A.U. *p<0.05, **p<0.01 and ***p<0.005, relative to D0). GAPDH was used as housekeeping gene. (B) Immunodetection of ADRP1 (green) and PLIN1 (red) in adipocytes (H9) on D25. Nuclei were stained with DAPI (blue). Higher magnification shown in merged image. Bar: 100 µm. (C) Representation of LipidTOX immunodetection at day 0, 8, 12, 20 and 25 of differentiation. Bar: 100 µm. (D) LipidTOX immunodetection quantification reveals increased levels of integrated intensity (left), mean lipid area per cell (middle) and percentage of LipidTOX positive cells (right) over the course of differentiation. Bar chart of integrated intensities represents mean ± SEM measured in the cytoplasm of individual cells. Other bar charts represent the mean ± SEM of n=5 technical replicates (*p<0.05, **p<0.01, ***p<0.001 and ****p<0.0001 compared to D0, Kruskal-Wallis).

A key feature of adipocytes is their responsiveness to insulin. Assessment of the insulin sensitivity of our cells at D25, by measuring phosphorylation of AKT at serine 473 following a 10 min stimulation with 100 nM insulin revealed an increase of AKT phosphorylation in insulin-treated vs untreated BAs (Fig.2D). Analysis of phosphorylation and total protein expression of other members of the insulin signalling pathway i.e., PTyr-IRS1/IRS1, P-P70S6K/P70S6K (Fig.2D), further validated the insulin sensitivity of our hPSC-derived BAs.

### Human PSC-derived brown adipocytes express thermogenic markers

Sustained induction of *C/EBP*β was reminiscent of a brown-like adipogenic program ^29^, and other BA markers further confirmed this. hPSC-derived BAs displayed a canonical thermogenic signature defined by *PRDM16, UCP1*, and *ZIC1* (Fig.4A), all reported to be classical BA markers ^30^. This signature was induced between D20 and D25 of differentiation and confirmed by immunofluorescence, showing lipid-loaded cells were positive for PRDM16 and ZIC1. Moreover, the same lipid-loaden cells co-expressed the mitochondrial markers UCP1 and COXII (Fig.4B-C and Fig.S4B-C). The expression of UCP1 in our cells during differentiation was confirmed by western blotting. UCP1 was expressed in hPSC-derived adipocytes at D20 and D25 at levels that were comparable to mature brown adipocytes from the PAZ6 BA cell line (Fig.4D and Fig.S4C). In addition to UCP1, we also detected high levels of PPARα, DIO2, and PGC1α (Fig.4D) at D20 and D25. Furthermore, these cells also expressed *ADRB3* (Fig.4A), which encodes the β3 adrenergic receptor. The β3 adrenergic receptor is the primary β-adrenergic receptor responsible for the activation of mature BAs *in vivo* ^*31*^. Complementary to this candidate approach, we also applied pathway analysis to RNAseq data and demonstrated that the transcriptome profile of differentiating cells at D25, correlated more strongly with the GO annotation “brown fat cell differentiation” than the one from PAZ6 human BAs (Fig.4E and Fig.S5A-B).

**Figure 4.**
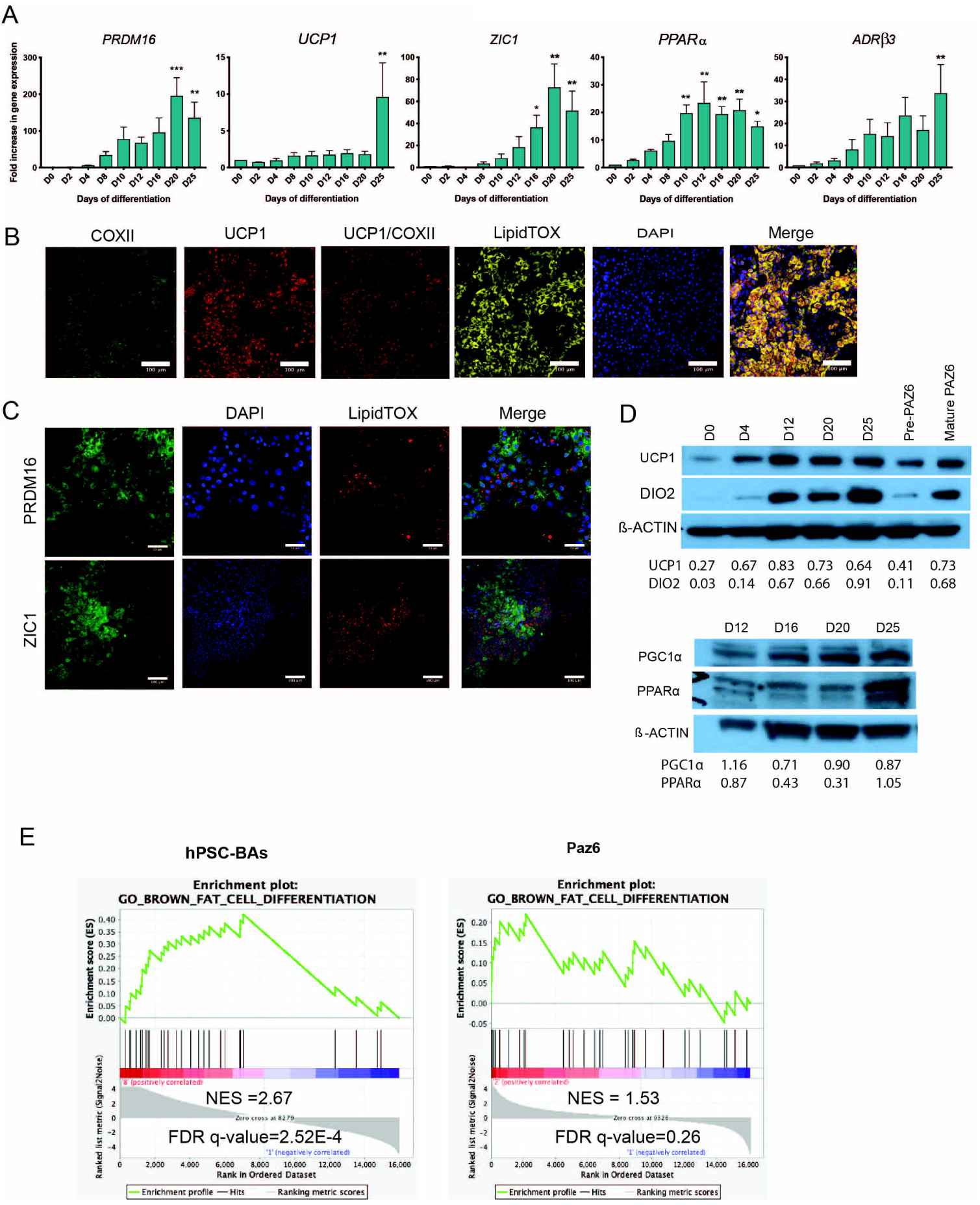
Brown adipocytes thermogenic markers. (A) Time-course mRNA expression of indicated brown adipocyte markers during differentiation of H9 (n ≥ 3 experiments; mean ± SEM A.U. *p<0.05 *p<0.05, **p<0.01 and ***p<0,005 relative to D0). GAPDH was used as housekeeping gene. (B) Immunodetection of COXII (Green) and UCP1 (red) in lipid-containing (LipidTOX - grey) adipocytes (H9) on D25. Nuclei were stained with DAPI. Bars: 100 µm. (C) Immunodetection of PRDM16 (up) and ZIC1 (down) (green) in lipid-containing (LipidTOX - red) adipocytes (H9) on D25. Nuclei were stained with DAPI. Bars: 100 µm. (D) Detection of UCP1 and DIO2 in H9-derived brown adipocytes on D0, D4, D12, D20 and D25 and in brown adipocytes PAZ6-cells in progenitors and mature stages by western blotting. Detection of PPARα and PGC1α in H9-derived brown adipocytes on D12, D16, D20 and D25 and in mature brown adipocytes PAZ6-cells by western blotting. β-ACTIN was used as loading control. Western blot quantification is shown underneath the WB image. (E) Gene Set Enrichment Analysis of H9-derived adipose cells at D25 *vs* D0 and PAZ6 human brown adipocytes D14 *vs* D0 with GO dataset (“brown fat cell differentiation”, GO:0050873), (n = 3 independent experiments).

### The transcriptional regulation of human PSC-derived brown adipocytes

To further strengthen the previous data, we first identified a nine transcriptional regulator signature activated during the differentiation of PAZ6 BAT cells. From these nine transcription factors, *PPARG and SOX13* regulate brown adipogenesis (Fig. 5, left) ^1,32^. Furthermore, eight of them were activated during specific steps of the differentiation of hESC cells into brown adipocytes and also all nine were activated during iPSC to BA differentiation (Fig. 5). Only one of them, *FOXO3*, was activated when the same ES cell line was differentiated to skeletal muscle (Fig.5, right) ^33^. Given the similarities in terms of developmental origin between skeletal muscle and brown adipose tissue, this analysis confirmed the specificity of our protocol for brown adipose tissue.

**Figure 5.**
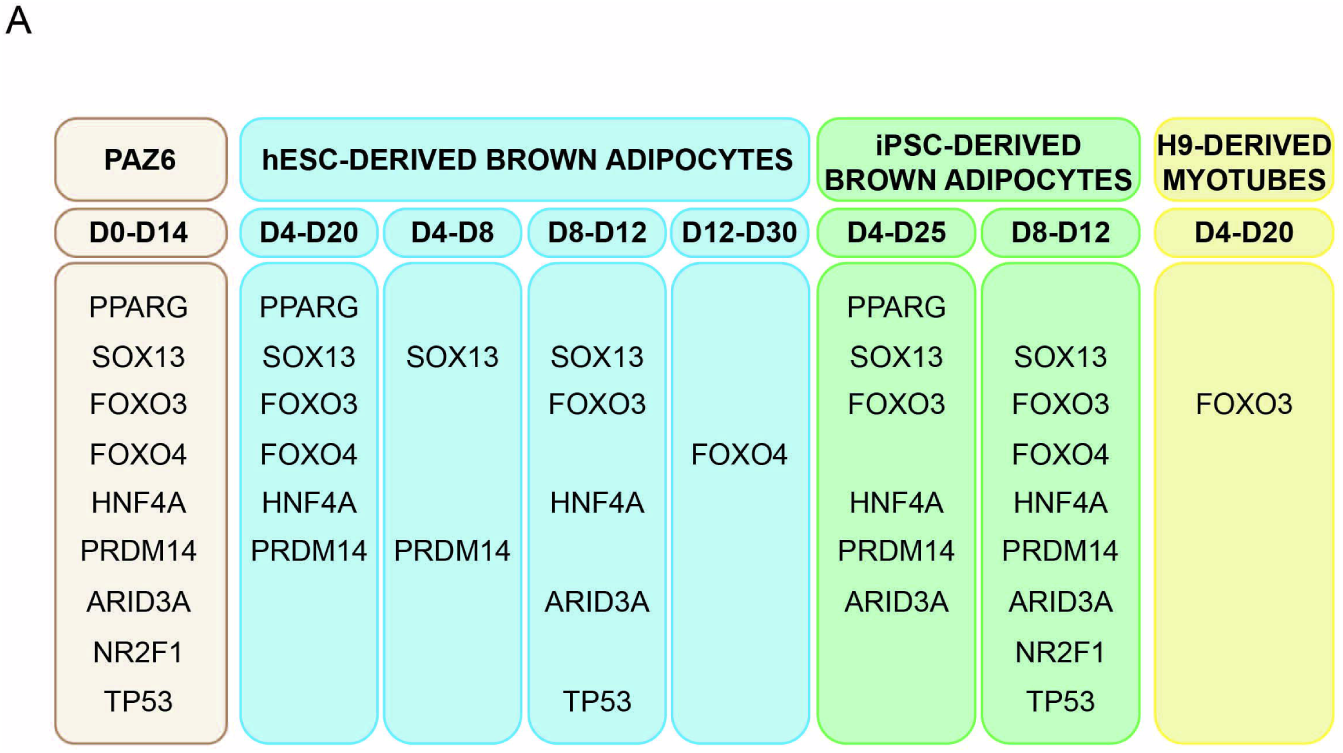
Transcriptional regulators of brown adipose tissue differentiation. (A) Identification of the transcriptional regulators activated in differentiating PAZ6 cells using the VIPER algorithm (left) and their correspondence in the indicated steps of hESC-(in blue) and iPSC-derived brown adipocytes (in green). HESC-derived myotubes have been included as a negative control.

Altogether, these results outline the specificity of our protocol for human PSC to BA differentiation in generating BAs with all the hallmarks of primary BAs isolated from mice and humans.

### hPSCs-derived brown adipocytes are functional

A key functional feature of BAs is their capacity to activate the thermogenic program through either β-adrenergic signalling or thyroid hormone. We confirmed that our BA cell system was able to produce a remarkable functional human BA that responds to norepinephrine (NE) and mirabegron (MIRA), a β3-adrenergic agonist ^34^. NE and MIRA stimulation for 2h also increased glucose uptake (Fig.6A and Fig.S4D-E). Furthermore, immunoblot analysis showed that the treatment of our hPSC-BAs with NE for 6h increased UCP1 and DIO2 protein expression (Fig. 6B). However, overnight treatment with T3 did not further induce UCP1 or DIO2 expression. We also tested the ability of different β-adrenergic stimuli to increase intracellular cAMP levels. Firstly, we determined that the general cAMP stimulating agents isoproterenol (IPR) and forskolin (FSK) were able to increase cAMP, indicating that the cells had functional adenylate cyclase as well as phosphodiesterases ^35^. More specifically, we demonstrated the responsiveness of cAMP levels to canonical activators of BAT, by treating cells with mirabegron (Fig.6C and Fig.S4F). In this context, β3 activation led to functional down-stream readouts of brown adipocyte activity. Mirabegron reduced lipid droplet (LDs) size and increased mitochondrial number (Fig 6D), both known indicators of BAT activation *in vitro* and *in vivo*. Oxygen consumption analysis of human PSC-derived BAs treated with mirabegron showed an increase in the basal respiration and ATP production *vs*. untreated controls (Fig.6E). The respiration profile of our cells is similar to that of mature murine brown adipocytes (Fig.S4G). Thus, this as a whole indicates that hPSC-derived adipocytes obtained from this protocol are fully functional BAs.

**Figure 6.**
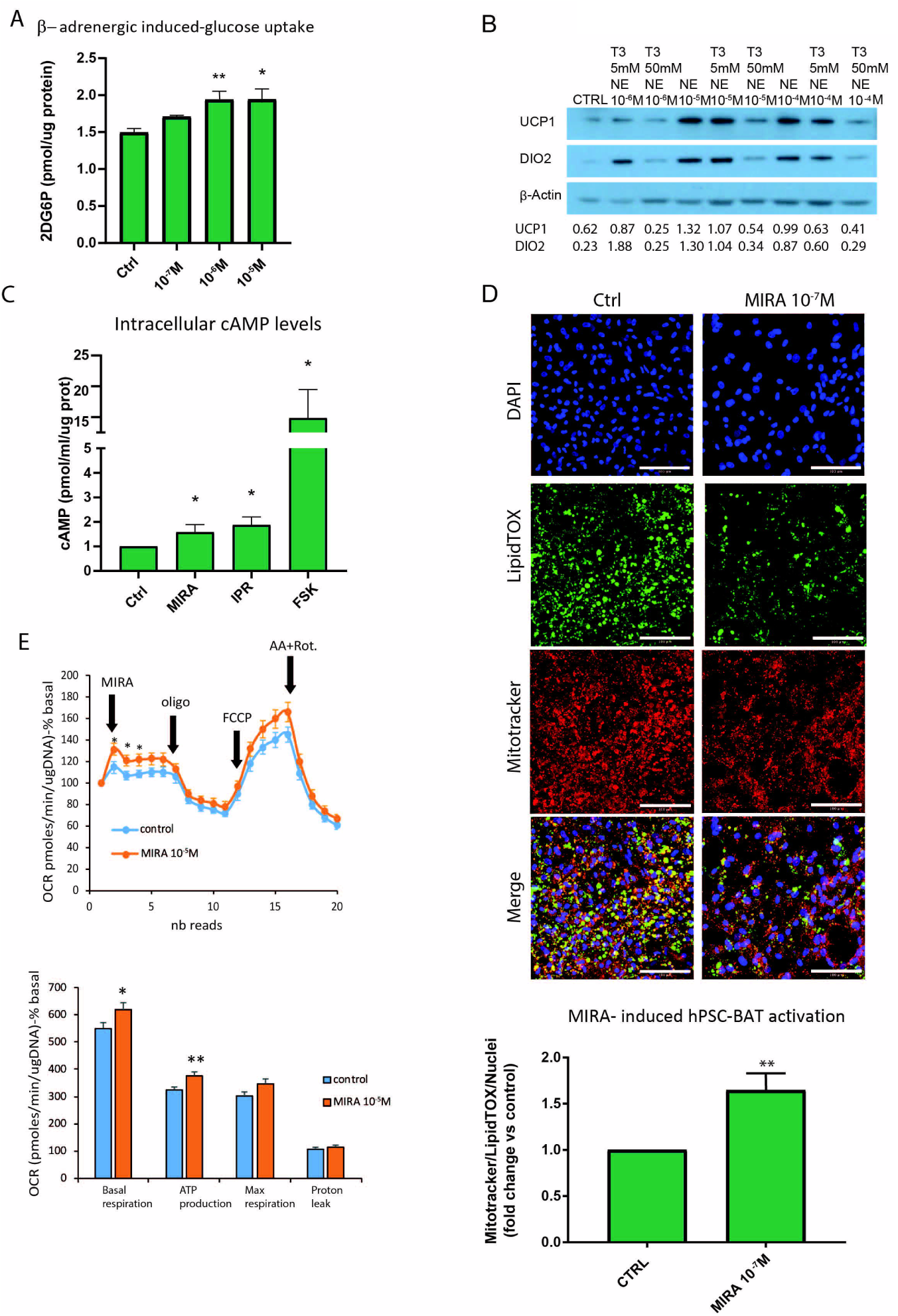
Brown adipocytes functional features. (A) β-adrenergic induced glucose uptake was evaluated with the Glucose uptake assay kit (Abcam). At D25, the cells were treated with different concentrations of NE (as indicated in the figure panel) (mean ± SEM n ≥ 3 experiments; **p<0.005 and *p<0.05 relative to control; Kruskal-Wallis test). (B) UCP1 and DIO2 western blot analysis at D25 after incubation at different concentrations of NE. β-ACTIN was used as loading control. (C) cAMP response to NE, IPR and FSK, all used at a concentration of 10^−5^M, was evaluated with cAMP parameter assay kit (R&D) (mean ± SEM n = 3 experiments;*p<0.05 relative to control; Kruskal-Wallis test). (D) Upper panel: Evaluation of lipid-droplet size and activation in a basal state (left panel) and after treatment with MIRA (right panel) using Mitotracker (red) and LipidTOX (green) in differentiated adipocytes at D25 using the same confocal settings. Nuclei were stained with DAPI. Bars: 100 µm); Lower panel: Calculation of the Mitracker on LipidTOX fluorescence ratio on the number of nuclei giving an indirect measure of the number of activated adipocytes in basal versus treated conditions. (Mean ± SEM n = 3 experiments, 5 fields per samples; **p<0.005 relative to control; Kruskal-Wallis test.) E) Seahorse XF analyser profile and quantitative summary of human PSC-derived BAT stimulated with or without 10^−5^M mirabegron (MIRA), following by treatment with 2µM oligomycin (oligo), 5µM FCCP and 1µM antimycin/rotenone (AA+Rot). (Mean ± SEM n = 22-25 wells;*p<0.05 relative to control; Student’s t-test).

### Our hPSC differentiation system is a useful tool for the temporal analysis of human brown adipocyte differentiation

As the cellular model is supposed to recapitulate the developmental steps undergone during brown adipogenesis, we validated the temporal expression of some of the known brown adipose tissue regulators, markers, and other functionally relevant proteins in humans ^36,37^. This analysis unveiled that *C/EBP*α, *C/EBP*β, and *PRDM16*, for example, are already upregulated in the adipose progenitor state (D8) Another transcriptional regulator such as *CIDEC* and *PPARγ* are, as could be anticipated, are specifically upregulated later in the differentiation (Fig. 7, upper row). Processes such as mitochondrial enrichment and upregulation of fatty acid oxidation enzymes occurred predominantly in the intermediate steps (BAs progenitors/preadipocytes) (Fig.7). This model is also useful in identifying unexpected comparative biology. For example, when comparing to mouse data, *ADRB3* and the hormone-sensitive lipase (*LIPE*) were found to be already expressed from the first precursor stage, rather than appearing as late-stage differentiation markers (Fig. 7). While needing confirmation, by allowing us to parse different developmental stages, our system suggests there may be significant differences between mice and humans in terms of the temporal expression of BA genes. Overall, these results indicate that our step-wise, chemically defined hPSC-to-BA culture system represents a potentially powerful tool to gain new insights regarding human BAs development and differentiation that could prove essential to make BAT a proper therapeutic target in humans.

**Figure 7.**
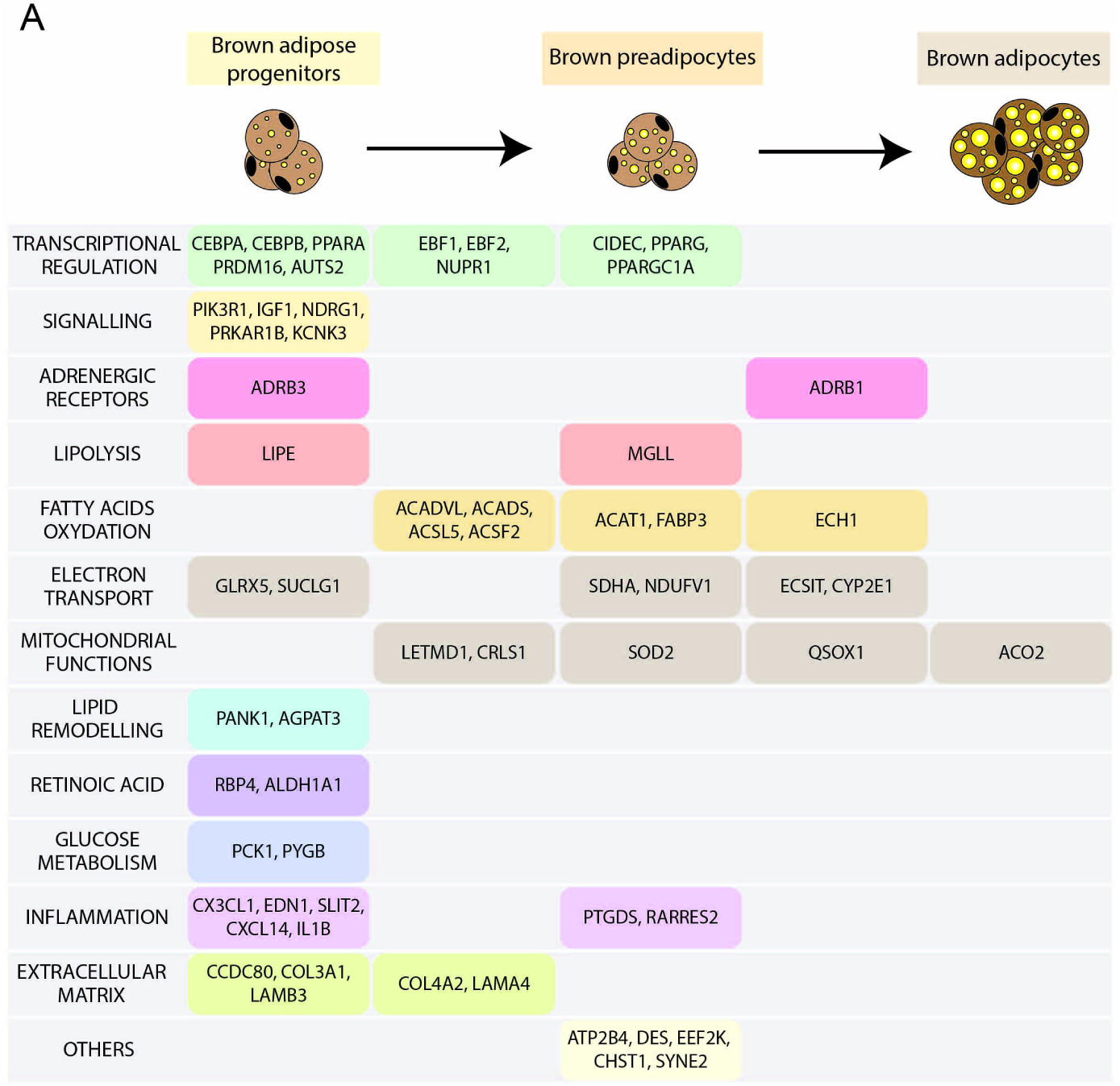
Temporal expression of human BAT markers and functionally relevant proteins. (A) Analysis of the first developmental step in which the expression of each of the indicated genes is upregulated based on the RNA-seq data. “Brown adipose progenitors” corresponds to D8, “brown preadipocytes” to D12 and “brown adipocytes” to D30. The data shown correspond to both H9 and Kolf-C2.

## Discussion

In this manuscript, we have elucidated a step by step chemically defined method to differentiate hPSCs into mature BAs. The importance of this method is that it takes pluripotent stem cells through the journey defined by specific developmental stages, closely mimicking the normal developmental program of BAT *in vivo*. Previously published protocols for the differentiation of hPSCs to BAs use either forced ectopic gene expression of the critical final transcription factors and/or employ embryoid body formation and MSC derivation ^15-19^. Although successful in generating mature BAT, these cellular models fail to capture the sequence of significant steps of human BAT formation, bypassing critical intermediate cellular identities in its development. Our unique vision is that if the ultimate goal is to increase BAT mass in the obese and diabetic population, having access to a clear and detailed characterisation of the critical intermediate cell stages connecting PSCs and BAs, is a unique opportunity to rescue the patient’s precursors. Targeting intermediate stages with drugs may boost the endogenous capacity for BAT mass formation.

For this vision to succeed, the main constrain is that the developmental origins of human BAs are not well defined. Most of the available information related to BAT development comes from murine studies ^38^ that have provided the primary evidence that BAs derive from the mesodermal germ layer ^39^, and more specifically from paraxial mesodermal progenitors ^22^. Our first question was whether humans might share similar stages for which we optimised a protocol recapitulating the different signals PSCs see in their journey to become a mature BA. For this, we took human PSCs through the same sequential cell-fate specification seen in mice. The fact that this road ultimately develops BA indicates that both species have a high degree of transcriptional similarity, as indicated by the clustering observed between primary human brown adipocytes and established cell lines. Thus we conclude that there is enough similarity in the differentiation pathway between rodents and humans to take advantage of the murine information.

We first induced PSCs toward early mesoderm by using a combination of insulin, FGF2, and the GSK3 inhibitor, Chiron, for two days. This step was followed by two days of insulin, FGF2 and RA, to induce the formation of paraxial mesodermal-like precursors. The rationale behind these specific combinations and sequence of compounds for these particular time frames was to mimic the intrinsic cues that guide the differentiation towards paraxial mesoderm typically observed during the development of the vertebrate embryo *in vivo* ^40^, also accounting for our previous work on PSCs ^24^. Our protocol succeeded in generating *TBOX*^*+*^ early mesoderm progenitors, which subsequently developed into paraxial mesoderm precursors, expressing *MYF5* and *PDGFR*α. Whereas TBOX (Brachyury) is a well-known pan mesodermal marker ^24,41,42^, the induction of *MYF5* and *PDGFR*α confirmed the relevance of paraxial mesoderm genes ^25^. The specific cellular identity was confirmed by transcriptomics. While these results provided firm evidence that we had generated paraxial mesoderm, the question as to whether these paraxial mesoderm cells would be competent to form BAs had to be elucidated.

In the next stage of our protocol, we sought to drive paraxial mesoderm toward adipogenic precursors. For this, we optimised a cocktail including LDN, a BMP signalling inhibitor, and Chiron, a GSK3 antagonist. LDN and Chiron are known to drive paraxial mesoderm precursors towards presomitic mesoderm-derived lineages such as muscle and BAT ^24,43,44^. As sought, this treatment induced the expression of *PAX3* and *EBF2*, two brown cell lineage progenitor markers ^17,45^. Moreover, GSEA analysis further confirmed that the cells generated from the paraxial mesoderm using this protocol were committed to the brown lineage.

At this point, the exposure of our adipogenic precursor cells to a classic adipogenic induction cocktail drove them toward terminally differentiated BAs. As they differentiated, our cells increased lipid accumulation and expressed mature adipogenic genes such as *PLIN1* and also thermogenic genes, including *UCP1, DIO2*, and *ADR*β*3*. Moreover, these hPSC-derived BAs expressed a range of transcription factors found in primary mature human brown adipocytes, further validating our cellular model.

The next question was whether we were making functionally competent BAs. We demonstrated that in response to NE, these cells increased glucose uptake ^46^, elevated cAMP levels in response to β3-agonist treatment ^47^, NE-induced expression of UCP1 and DIO2 ^48^ as well as increased basal respiration after treatment with a β3-agonist ^34^ all typical characteristics of a BA cell.

In summary, here we have developed a step by step, robust protocol that recapitulates the developmental stages transforming hPSCs into functional BAs following a rationally designed developmental program road map. This is a unique tool to identify essential key regulatory factors operating at specific stages governing the stage-specific decisions required to guide the PSC towards becoming a fully functional BAs. Our protocol avoids the issues caused by the shortcut resulting from engineered ectopic gene expression systems, where changes in expression is driven by the intense action of the transcription factors introduced. Furthermore, the scalability of our protocol, we think this is one of its main assets, as it provides a reproducible resource to gain new insights into human BAT formation and physiology suitable for screenings at different developmental stages. Ultimately, we envision that the knowledge generated from studies using this source of inexhaustible human brown fat may allow the development of new safe treatments for obesity and diabetes.

## Materials and methods

### Cell culture and maintenance

Two hPSCs lines were employed. The hESC line H9 (WA09, WiCell, Madison, WI) was maintained in Essential 8(tm) (E8) medium (Gibco) on Vitronectin XF(tm) (Stemcell technologies) coated (1:1000 dilution) tissue culture-treated dishes and passaged mechanically using PBS-EDTA. The hiPSC line KOLF2-C1, a subclone of the hiPSC KOLF2 cell line (HPSI0114i-kolf_2, Human Induced Pluripotent Stem Cell Initiative (HipSCi), http://www.hipsci.org) were maintained in TeSR(tm)-E8(tm) (Stemcell Technologies) on Synthemax® II-SC Substrate (10µg/mL) (Sigma-Aldrich) and passaged mechanically using PBS-EDTA or Gentle Cell Dissociation Reagent (Stemcell Technologies). The human immortalised brown adipocyte cell line Paz6 was cultured as described previously 12. Mouse adipocytes were cultured as previously described ^49^.

### Cell differentiation

For differentiation, pluripotent cells were plated into Matrigel-coated 12-well plates and induced when 70% confluent. At days 0-4, chemically defined medium (CDM; BSA and insulin-free, prepared by Cellular Generation and Phenotyping (CGaP), Wellcome Sanger Institute (Table S1), Hinxton UK was used. From days 6-30, complete medium (DMEM-F12 Ham) (see Table.S2) was used. For functional analyses, cells were plated onto Matrigel-coated glass-bottomed 96-well plates (Eppendorf) at day 4 of differentiation; cells were detached using TrypLE (Gibco) and plated at single-cell suspension (one 12-well plate for two 96-well plates).

### RNA extraction

Cells were washed three times with PBS and lysed in RLT buffer containing 1% β-ME. Cell lysates were passed five times through a 23G needle before proceeding to RNA extraction using RNeasy Qiagen Kit (Qiagen) following the manufacturer’s instructions. RNA concentration was quantified using an Epoch(tm) 2 microplate reader.

### Realtime qPCR

RT-qPCR assessed mRNA levels of genes of interest. cDNA was generated from 500ng isolated RNA using M-MLV reverse transcriptase (Promega) and diluted 1:5 for use in 12µL qPCR reactions using SYBR® Green PCR master mix (Applied Bioscience) run on the Applied Biosystems StepOnePlus(tm) system Applied Biosystem, Carlsbad, California). Expression values are normalised to GAPDH (See Table.S3).

### Total RNA library construction and RNAseq

RNA samples were quantified with QuantiFluor RNA System, 1ml from Promega UK Ltd using Mosquito LV liquid platform, Bravo WS and BMG FLUOstar Omega plate reader, and cherrypicked to 100ng / 50µl using Tecan liquid handling platform. Library construction (poly(A) pulldown, fragmentation, 1st, and 2nd strand synthesis, end prep, and ligation) was carried out using ‘NEB Ultra II RNA custom kit’ on an Agilent Bravo WS automation system.

PCR was set-up using KapaHiFi Hot start mix, and unique dual indexed tag barcodes on the Agilent Bravo WS automation system. Post PCR, the plate was purified using Agencourt AMPure XP SPRI beads on Caliper Zephyr liquid handling platform. Libraries were quantified with Biotium Accuclear Ultra-high sensitivity dsDNA Quantitative kit using Mosquito LV liquid handling platform, Bravo WS and BMG FLUOstar Omega plate reader. Libraries pooled in equimolar amounts on a Beckman BioMek NX-8 liquid handling platform and pooled libraries were quantified on an Agilent bioanalyser. Libraries were normalised to 2.8nM. Samples were sequenced on a HiSeq 2500 platform.

### RNAseq data analysis

Fastq files were processed through a customised pipeline. The adapters were hard clipped before alignment through Cutadapt v. 2.3. The alignment was performed using STAR v. 2.5 on the GRCh.38. The reads per gene were counted, relying on feature Counts (Subread v. 1.6.4). Differential transcriptome analysis was performed using DESeq2, version 1.20.0.

### Analyses of the transcriptional regulators

The inference of the upstream transcriptional regulators was performed with VIPER (Virtual Inference of Protein-activity by Enriched Regulon analysis) ^50^. The algorithm provides a Normalised Enrichment Score (NES), which determines the activation/inhibition status of the transcription factor of interest, based on the observed differential expression of their known gene targets. The TF – target gene interactions network was obtained by DoRothEA ^51^. The outcome of VIPER analysis is a positive NES for activated and negative NES for inhibited TFs, respectively. Only transcription factors with a p-value lower than 0.05 were considered as statistically significant. RNA-seq data for H9-derived myotubes were obtained from the public repository GEO (GSE121154) and analysed as above.

### GSEA analysis

Gene Set Enrichment Analysis (GSEA; www.broadinstitute.org/GSEA) was conducted on pre-ranked and non-pre-ranked lists of genes. The ranking was computed according to the differential transcriptome analysis mentioned above. In particular, we calculated the -log_10_ of the adjusted p-value referring to the Wald test used for determining the significance of the differential transcriptome expression and we assigned to this value the sign of the fold change. The GSEA analysis was performed using 1000 gene set permutations, and no collapsing. Gene set sizes were selected to be 15-500, classic enrichment and meandiv normalisation mode. The databases used for analyses were: C5.all.V7.0 for D4 and D25 *vs* D0 is and C2.all.V7.0 for D12 *vs* D0. GSEA analysis of CPM non pre-ranked list were processed with the following parameters: the C5.bp.v7.0 database was used with 1000 gene set permutations, and no collapsing. Gene set sizes were selected to be 15-500, Signal2noise metrics for ranking genes, classic enrichment, and meandiv normalisation mode.

### Immunocytochemistry

For immunocytochemistry, cells were fixed in 4% PFA for 15 min at room temperature and blocked using 3% FFA-free bovine serum albumin (BSA) 0.1 % Triton X100 or saponin in PBS. Primary antibodies (Table S4) were diluted in PBS, 0.01 % Triton X100 or saponin, 1% FFA-free BSA. Secondary antibodies coupled to Alexa Fluor 488 and 546 or 627 (Raised in Donkey, Thermofisher Scientific) were diluted to 1/1000 in PBS, 0.01% Triton X100 or saponin, 1% FFA-free BSA. Adipocytes were incubated for 45min with HCS LipidTOX Neutral Green / Red or Deep Red neutral lipid stain (Thermofisher Scientific). After a 5min incubation with DAPI (SIGMA, 1/10000), cells were mounted in Fluoromount-G (Southern Biotech) (Table S4). Cell preparations were observed with a confocal microscope (Zeiss LSM 700). Immunocytochemical quantification was performed with the ICY image analysis software (http://icy.bioimageanalysis.org/).

### Immunoblot

Cells were lysed in cold RIPA buffer and homogenised by vortexing and running samples 5 times through a 23G needle. Samples were spun at maximum speed for 15 min at 4°C, and the supernatant collected. Protein was quantified using BIO-RAD DC(tm) protein assay (Biorad) following the manufacturer’s instructions. Samples of 20µg protein in RIPA and 1X loading buffer (10% β-Mercaptoethanol) were denatured at 95°C for 5min and run on NuPAGE(tm) 4-12% Bis-Tris Midi Protein Gels (Invitrogen) at 200V for 1h. Protein was transferred to PVDF membranes (Invitrogen) using iBlot(tm) dry transfer (Thermofisher Scientific). To block unspecific binding of antibodies, membranes were incubated in PBS-T 5% milk for 1h at RT. Primary antibodies were diluted in PBS-T 3% BSA (see Table.S3), and secondary antibodies were diluted in PBS-T 5% milk (see Table.S4).

### Insulin sensitivity

Cells were treated with 100nM of insulin for 10 min, at day 25-30 of differentiation, after an o/n incubation in complete medium without serum. The insulin sensitivity of the cells was assessed by measuring the levels of p-AKT, tot AKT, p-IRS1, tot-IRS1, p-P70S6K, and tot-P70S6K. β-ACTIN was the loading control.

### Lipid quantification

Image visualisation was performed using Fiji software ^52^. Fluorescent images were analysed using CellProfiler 3.1.9 ^53^ using custom-built pipelines. After applying illumination correction, nuclei were identified using the Otsu two-class thresholding method on the DAPI image channel. Nuclei touching the border of the image were discarded. Subsequently, a cytoplasm mask was created by expanding the nuclei objects by 10 µm. After the exclusion of the nuclei from the cytoplasm, intensity features of the LipidTOX image channel were measured inside the cytoplasm area. Mean lipid area was obtained by measuring the area occupied by lipids normalised to the number of nuclei. The percentage of LipidTOX positive cells was calculated by setting a fixed intensity threshold.

### Seahorse oxygen consumption measurements

Cells were differentiated in 24-well Seahorse V17 culture plate for 25 days. Before oxygen consumption rate (OCR) assay, complete medium was replaced with Seahorse medium without serum and cytokines. With a Seahorse XF24 analyser, OCR was measured with small molecule inhibitors added through the injection ports. The following concentrations of activators and inhibitors were used: 100, 10 and 1µM mirabegron, 2µM oligomycin, 5µM FCCP and 1µM each of antimycin A and rotenone. Basal, uncoupled, and maximal respiration rates were calculated upon the subtraction of the non-mitochondrial oxygen consumption obtained at the end of each assay by the addition of antimycin A and rotenone. The values obtained were normalised to total mg DNA per well as measured by Quant-iT(tm) PicoGreen(tm) dsDNA Assay Kit (Invitrogen). For mouse adipocytes, the following concentrations of activators and inhibitors were used: 1µM oligomycin, 0.9µM FCCP, and 1µM each of antimycin A and rotenone.

### Glucose uptake

Norepinephrine (NE)-induced-glucose uptake was assayed according to the established protocol from a commercial glucose uptake kit (Abcam). In brief, at day 25-30 of differentiation, human PSC-derived BAs seeded in 12-well plates were fasted overnight in complete medium without serum. The next day cells were treated with vehicle or indicated concentration of NE. After 2h of incubation, cells were washed three times with cold PBS and lysed with extraction buffer, frozen at –80□°C for 10□min and heated at 85□°C for 40□min. After cooling on ice for 5□min, the lysates were neutralised by adding neutralisation buffer and centrifuged. The remaining lysate was then diluted with assay buffer. Finally, the colorimetric end product generation was set up by two amplification steps according to the manufacturer’s instructions in the kit and then detected at 412□nm using a Spark microplate reader (Tecan).

### cAMP measurements

The cAMP assay was performed using the cAMP ParameterAssay Kit (R&D Systems, Minneapolis, MN, USA). In brief, at day 25-30 of differentiation, hPSC-derived BAs seeded in 12-well plates were fasted overnight in complete medium without serum. The next day cells were treated with vehicle or indicated concentration of NE, mirabegron, isoproterenol, and forskolin for 2h. Cells were washed three times in cold PBS, resuspended in cell lysis buffer 5 (diluted 1:5)*, and frozen at ≤ -20 °C. Then the cells were thawed with gentle mixing. The freeze/thaw cycle was repeated once. Samples were then spun at 600 x g for 10 min at 2-8 °C to remove cellular debris, and the supernatant was stored at ≤ -20 °C. The assay was carried out following the manufacturer instructions (R&D Systems, Minneapolis, MN, USA).

### Statistical analysis

All analyses were performed with Graphpad Prism Software. After checking the normality of data obtained in each experiment, the appropriate statistical test was applied to calculate significance values between data sets. The following statistical analyses were used to calculate *P*-values: Ordinary one-way Anova and Kruskal– Wallis. Holm-Sidak’s and Dunn’s post-hoc tests were performed for multiple comparisons to reduce errors in Ordinary one-way Anova and Kruskal–Wallis analyses, respectively.

## Data availability

RNA-seq data have been deposited in GEO…

## Conflict of interest

The authors declare no conflict of interest.

## Author’s contribution

AVP and SC and BR conceived the original hypothesis. SC and ACG designed and performed *in vitro* experiments, part of the bioinformatics analysis and wrote the manuscript. MB, IS, KL and FH, performed *in vitro* experiments, discussed and edited the manuscript. DC performed the RNAseq bioinformatics analysis. SRF, IK and SA performed part of the bioinformatics analysis. SM contributed to design some of the experiments, AB and LV contributed by providing advice and some of the cell lines used in this work. DL and SE participated by guiding some of the *in vitro* experiments. AVP wrote the manuscript and is the guarantor of this work. All authors approved this publication.

## Acknowledgements

We thank the CGaP team at Sanger for their technical assistance, the Molecular cytogenetics Lab at Sanger for the karyotyping of the cells and the DNA pipeline at Sanger for running the RNAseq. The PAZ6 cell line was a kind gift from Dr Tarik Issad, Institute Cochin, and Paris, France. We thank Dr Sergio Rodriguez-Cuenca for his scientific and technical advice and Dr Sam Virtue for his technical advice and revision of the manuscript. This work was funded by the ERC Senior Investigator award (669879). LV lab is funded by the ERC advanced grant New-Chol, the Cambridge University Hospitals National Institute for Health Research Biomedical Research Centre, and a core support grant from the Wellcome and MRC to the Wellcome – Medical Research Council Cambridge Stem Cell Institute.

## Figure Legends

Supplementary information is available at Cell Research’s website.

## Graphical abstract

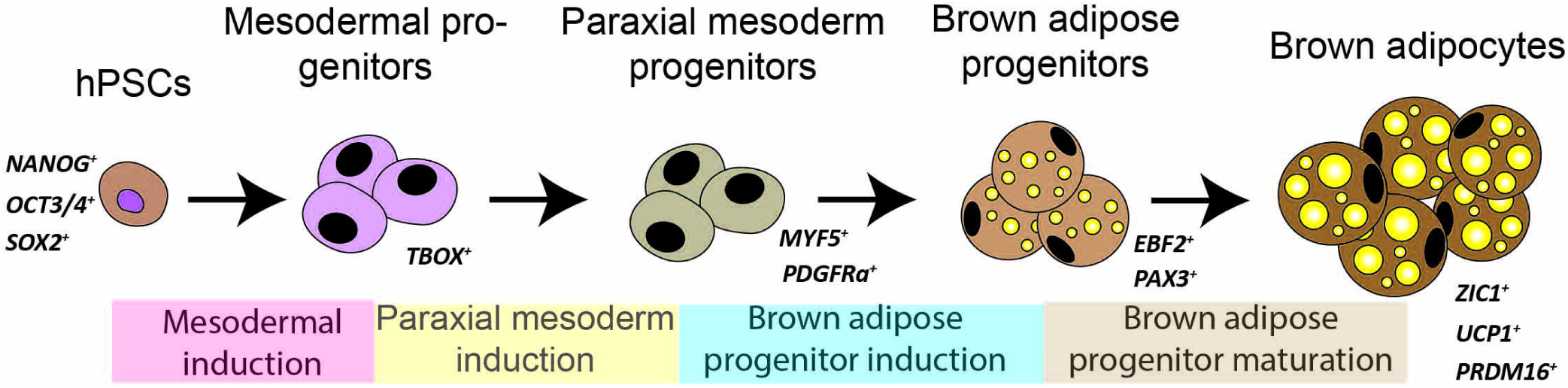

## Notes

### Competing Interest Statement

The authors have declared no competing interest.

