## supplementary information for "Unravelling the developmental roadmap towards human brown adipose tissue"

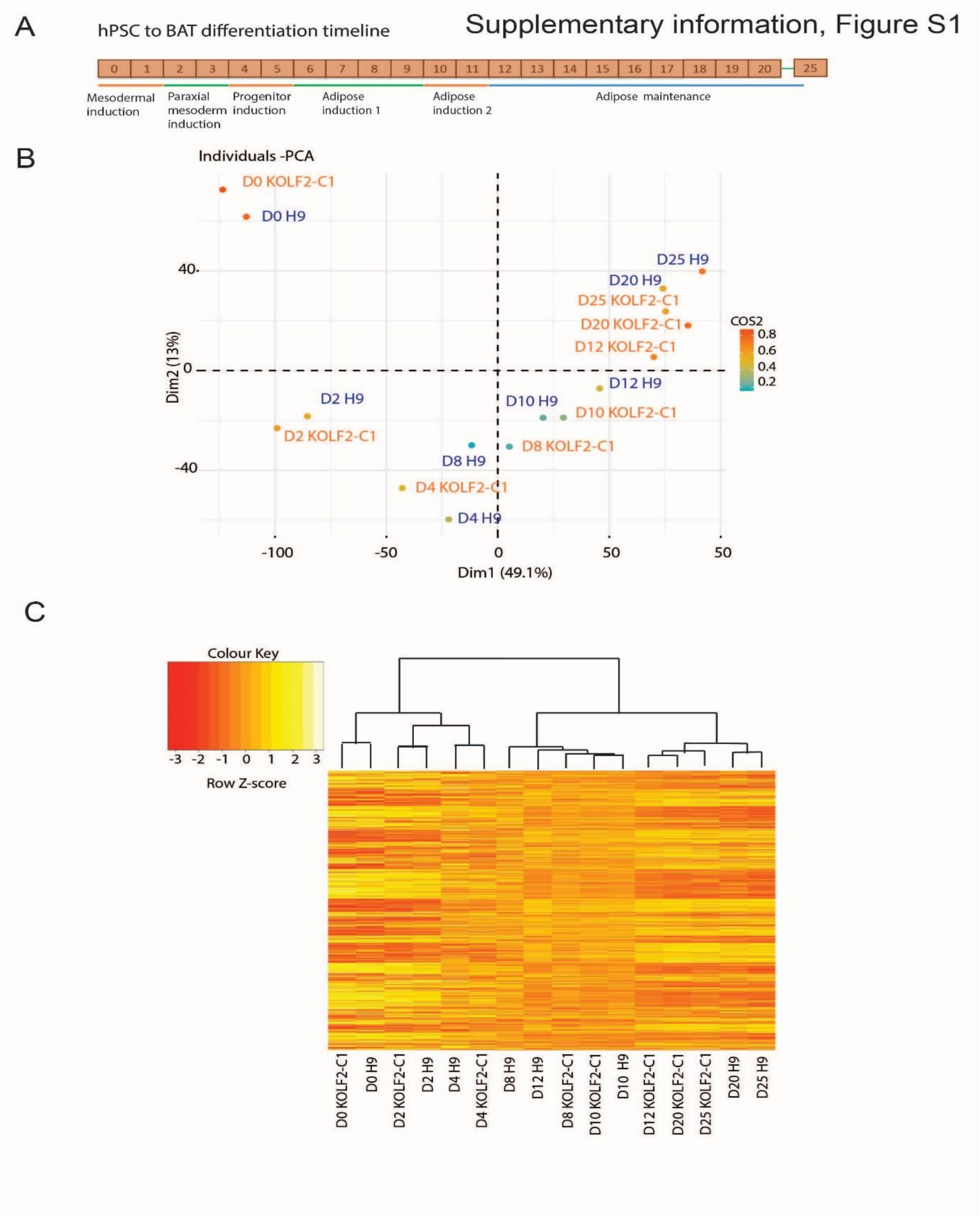


**Figure S1. PCA and clustering analysis of H9 and KOLF2-C1 cell lines.**

(A) Human PSCs to BAT differentiation protocol timeline.

(B) PCA plot of H9 human ES (here h9) and KOLF2-C1 (here Kolf2) hiPS cell lines differentiation into BAs RNAseq timepoints (D0, D2, D4, D8, D10, D12, D20 and D25). (H9, n=3 and KOLF2-C1 n=5 independent experiments).

(C) Heatmap of clustering analysis of H9 human ES and KOLF2-C1 hiPS cell lines differentiation into BAs. Unsupervised clustering of the D0, D2, D4, D8, D10, D12, D20 and D25 RNAseq timepoints. Upregulated genes are represented in yellow and downregulated clusters in red. (H9, n=3 and KOLF2-C1 n=5 independent experiments).

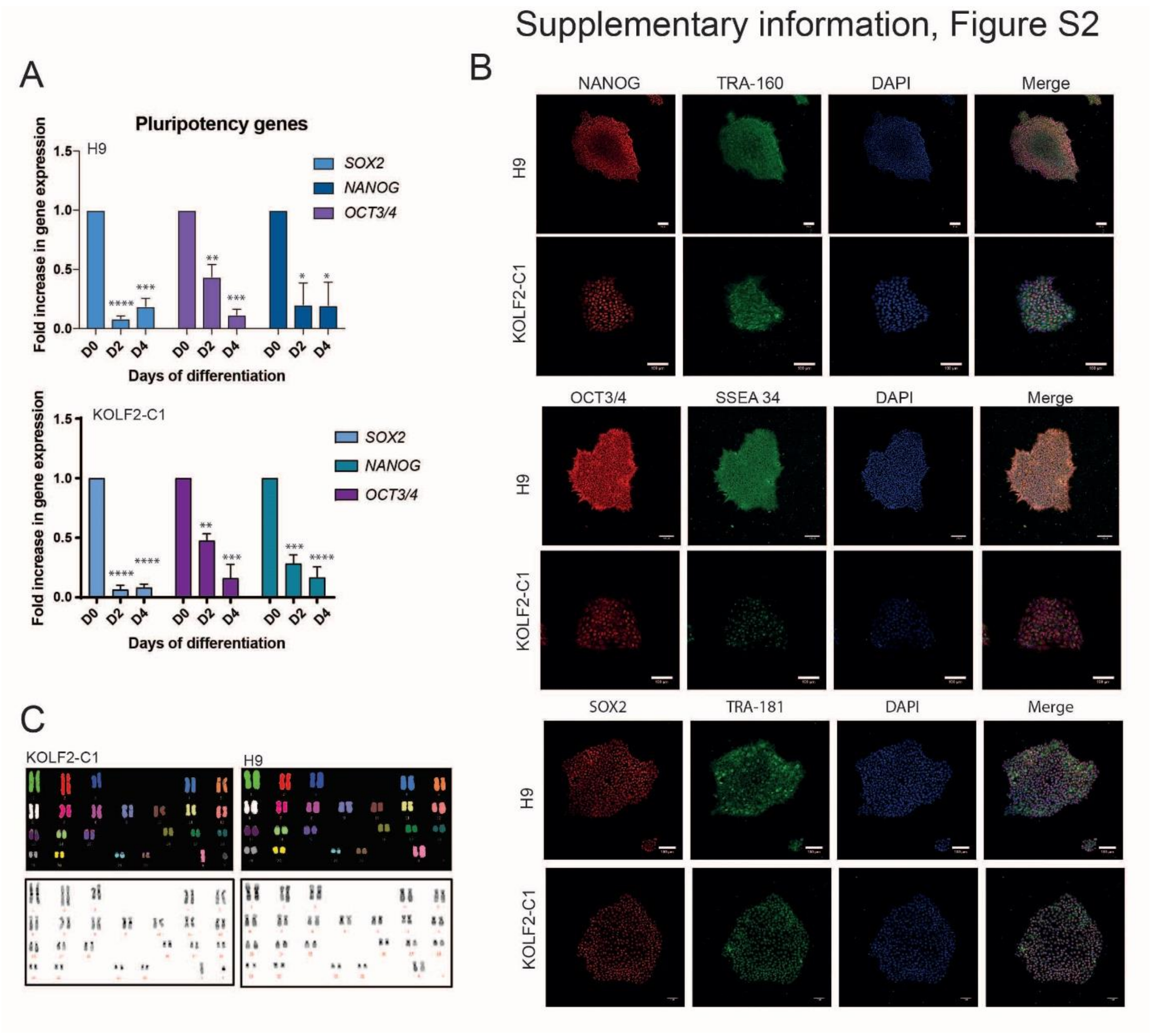


**Figure S2. Pluripotency analysis and karyotyping of H9 and KOLF2-C1 cell lines**

(A) RT-qPCR analysis of expression of pluripotency genes *OCT3/4, NANOG* and *SOX2* (mean ± SEM arbitrary units (A.U.) relative to D0; n ≥ 3 independent experiments; **** and ***p<0.0001, **p<0.005, *p<0.05 relative to D0) in H9 (upper panel) and in KOLF2-C1 (lower panel).

(B) Immunodetection of OCT3/4, NANOG and SOX2 (red) co-localised respectively with SSEA 3/4, TRA-1-60 and TRA-1-81 in pluripotent stem cells (H9 and KOLF2-C1) on D0. Nuclei were stained with DAPI. Bars: 100 µm

(C) Karyotyping analysis of H9 and KOLF2-C1 cells lines.

Supplementary information, Figure S3

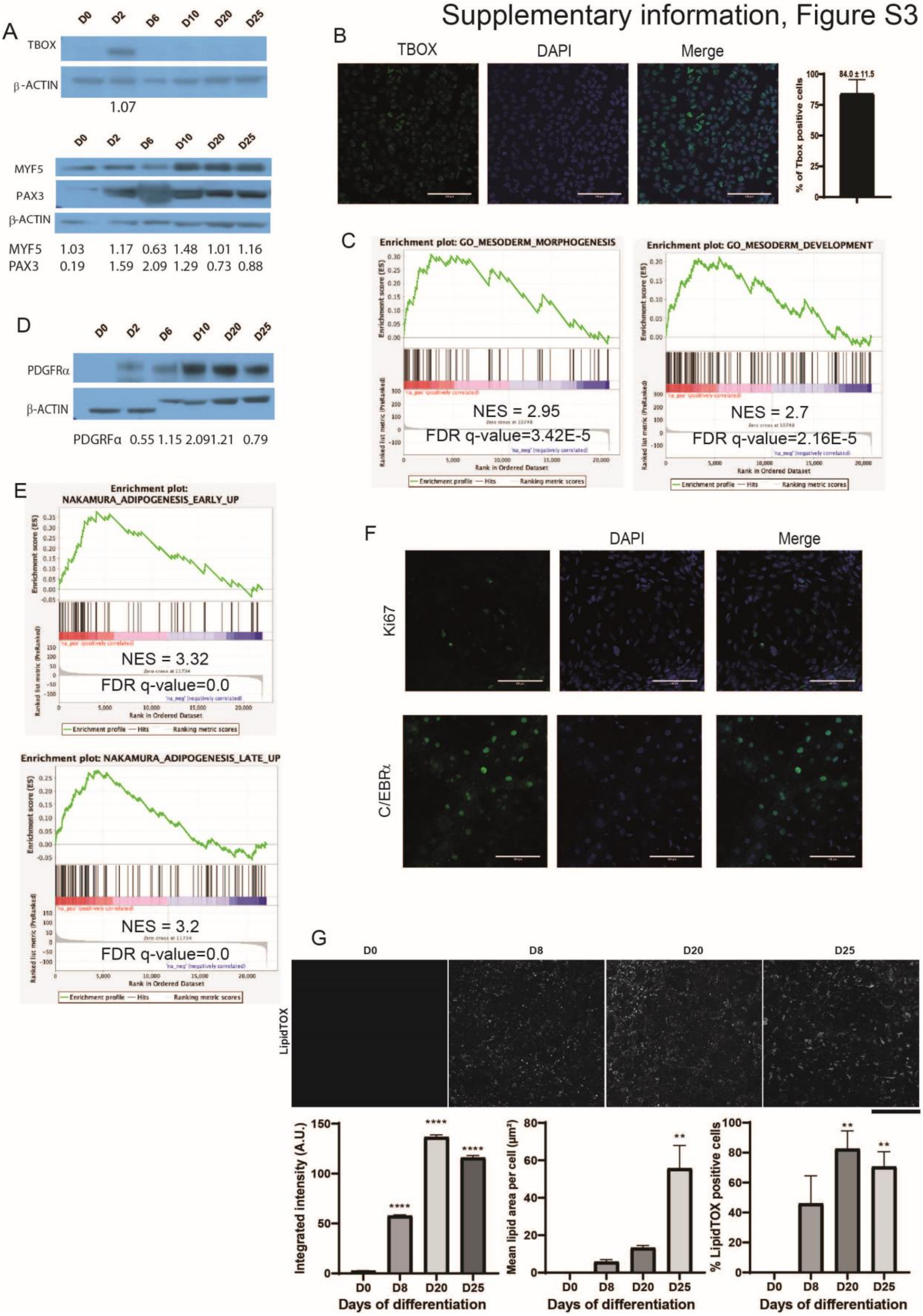

**Figure S3. Human iPSC-derived brown adipocytes progenitor molecular characterisation and lipid accumulation during dfifferentiation.**

(A) Detection of TBOX, MYF5, PAX3, in KOLF2-C1-derived brown adipocytes on D0, D2, D6, D10, D20 and D25 by western blotting. β-ACTIN was used as loading control. Western blot quantification is shown underneath the WB image.

(B)TBOX immunostaining of mesodermal progenitors at D2 (green). Nuclei were stained with DAPI. Bars: 100 µm. TBOX positive cells were quantified using CellProfiler (mean + SEM, n=3 technical replicates).

(C) Gene Set Enrichment Analysis of KOLF2-C1-derived cells on D4 *vs* D0 using GSEA (n=5 independent experiments) using the "mesoderm morphogenesis" GO:48332, "mesoderm development" GO:0007498 datasets.

(D) Detection of PDGFRα in KOLF2-C1-derived brown adipocytes on D0, D2, D6, D10, D20 and D25 by western blotting. β-ACTIN was used as loading control. Western blot quantification is shown underneath the WB image.

(E) Gene Set Enrichment Analysis of KOLF2-C1-derived adipose progenitors cells using published datasets ("Nakamura adipogenesis early up" and "Nakamura adipogenesis late up"), with early and late adipogenesis transcriptomic signatures on D12 *vs* D0, compared to human adult adipose stromal cell signature (n=5 independent experiments).

(F) Ki67 immunostaining of adipose progenitors at D12 (green). C/EBPα immunostaining of adipocytes at D25 (green). Nuclei were stained with DAPI (blue). Bars: 100 µm.

(G) LipidTOX immunodetection quantification reveals increased levels of integrated intensity (left), mean lipid area per cell (middle) and percentage of LipidTOX positive cells (right) over the course of differentiation. Bar chart of integrated intensities represents mean ± SEM measured in the cytoplasm of individual cells. Other bar charts represent the mean ± SEM of $n=5$ technical replicates (*$p<0.05$, **$p<0.01$, ***$p<0.001$ and ****$p<0.0001$ compared to D0, Kruskal-Wallis).

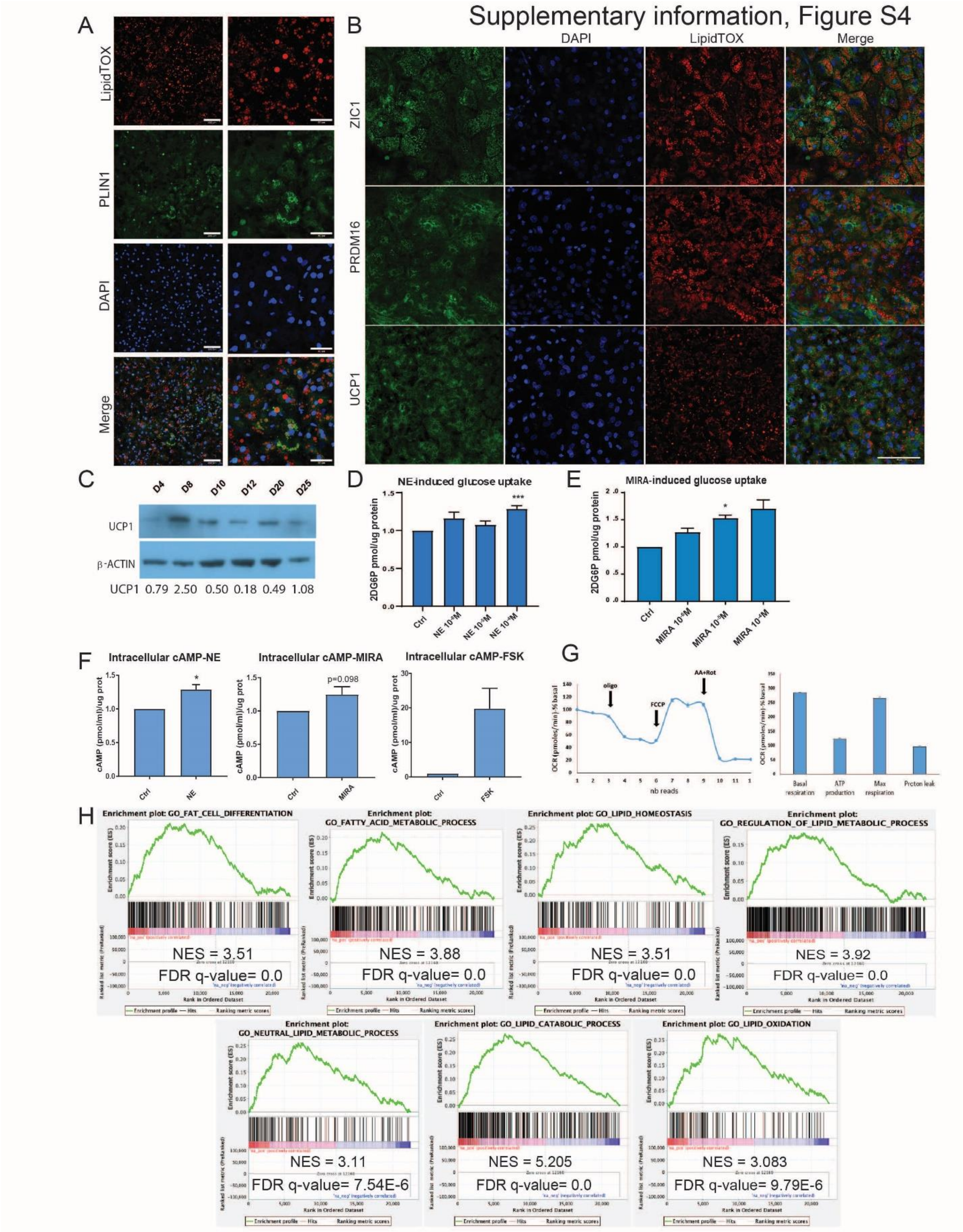


**Figure S4. Human iPS-derived brown adipocyte molecular characterisation and metabolic phenotyping.**

(A) Immunodetection of PLIN1 (green) in KOLF2-C1-derived lipid-containing adipocytes (LipidTOX-red) on D25. Nuclei were stained with DAPI. Bars: 100 µm.

(B) Immunodetection of ZIC1, PRDM16 and UCP1 (green) in lipid-containing (LipidTOX-red) adipocytes (KOLF2-C1) on D25. Nuclei were stained with DAPI. Bars: 100 µm.

(C) Detection of UCP1 in KOLF2-C1-derived brown adipocytes on D4, D8, D10, D20 and D25 by western blotting. β-ACTIN was used as loading control. Western blot quantification is shown underneath the WB image.

(D) NE- induced glucose uptake was evaluated with the Glucose uptake assay kit (Abcam). At D25, the cells were treated with different concentrations of NE (as indicated in the figure panel) (mean ± SEM n ≥ 3 experiments; ***p<0.001 and * relative to control; Kruskal-Wallis test).

(E) MIRA-induced glucose uptake was evaluated with the Glucose uptake assay kit (Abcam). At D25, the cells were treated with different concentrations of MIRA (as indicated in the figure panel) (mean ± SEM n= 3 wells; *p<0.05 relative to control; Kruskal-Wallis test).

(F) cAMP levels in response to NE, MIRA and FSK treatment, all used at a concentration of $10^{-5}$M, (mean ± SEM n = 2-4 experiments, *p<0.05 relative to control, Kruskal-Wallis test).

(G) Seahorse XF analyser profile and quantitative summary of mouse brown adipocytes following by treatment with 1µM oligomycin (oligo), 0.9µM FCCP and 1µM antimycin/rotenone (AA+Rot). (mean ± SEM n = 10 wells.

(H) Gene Set Enrichment Analysis of KOLF2-C1-derived adipose cells at D25 *vs* D0 with GO datasets (“fat cell differentiation” GO:0045444, “fatty acid metabolic process” GO:0006631, “lipid homeostasis” GO:0055088, “regulation of lipid metabolic process”

GO:0006629, “neutral lipid metabolic process” GO: 0006638, “lipid catabolic process” GO:0016042 and “lipid oxidation” GO: 0034440), (n = 5 independent experiments).

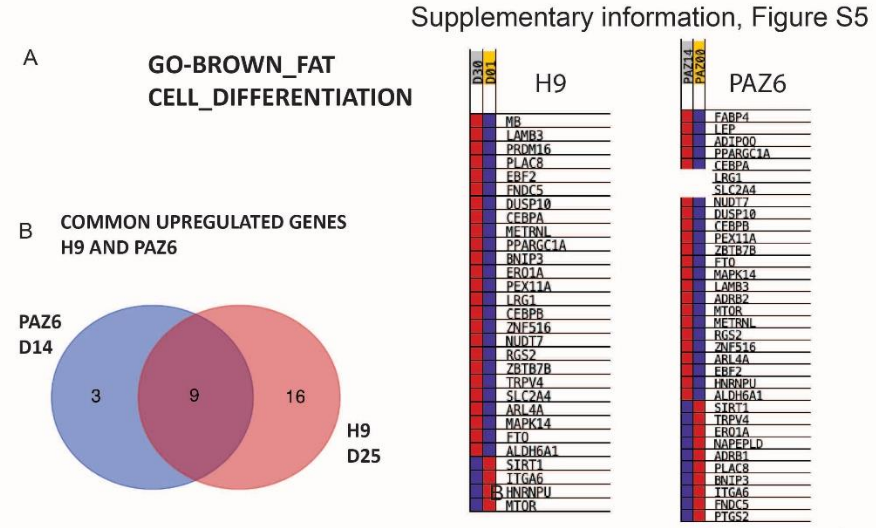


| Names | total | elements |
|---|---|---|
| H9 PAZ | 9 | DUSP10 NUDT7 ZBTB7B SLC2A4 LRG1 PEX11A CEBPB PPARGC1A CEBPA |
| PAZ | 3 | FABP4 LEP ADIPOQ |
| H9 | 16 | ARL4A PRDM16 MAPK14 ZNF516 TRPV4 METRNL RGS2 PLAC8 FNDC5 EBF2 MB LAMB3 ALDH6A1 ERO1A FTO BNIP3 |

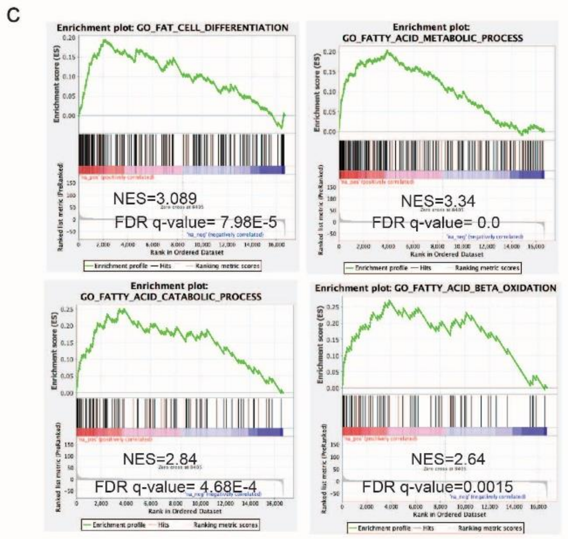


**Figure S5. Common upregulated genes shared between terminally differentiated human PSC-derived and PAZ6 brown adipocytes and GSEA analysis of mature PAZ6 cells.**

(A) Detailed heatmaps showing the gene expression of H9 at D25 and PAZ6 at D14 of differentiation for the dataset GO_Brown_fat_cell_cell differentiation.

(B) Common upregulated genes of H9 at D25 and PAZ6 at D14 of differentiation in the context of GO_Brown_fat_cell_cell differentiation.dataset.

(C) Gene Set Enrichment Analysis of PAZ6 human brown adipocytes at D14 *vs* D0 with GO datasets (“fat cell differentiation” GO:0045444, “fatty acid metabolic process” GO:0006631, “fatty acid catabolic process” GO:0006631 and “fatty acid beta oxidation GO:0006635), (n = 3 independent experiments).

**Table S1. Composition Chemically Defined Medium (CDM)**

| Compounds | Total volume/quantity (concentration) |
|---|---|
| F-12 Nut Mix (lnvitrogen 31765068) | 250ml (50%) |
| IMDM (lnvitrogen 21980065) | 250ml (50%) |
| HyClone BSA (GE Healthcare SH30574-02) | 2.5g (0.Smg/ml) |
| CD Lipid Concentrate (lnvitrogen 11905031 | 5ml (1%) |
| Insulin (Roche 1376497).Reconstituted in water at l0mg/ml | 350µ1 (7µg/ml) |
| Transferrin (Roche Sigma 10652202001) 30mg/ml | 250µ1 (15µg/ml) |
| Mono-Thioglycerol (Sigma M6145-25ml) 11.5M | 20µ1 (0.5mM) |

**Table S2. Cell culture medium composition at the different stages of differentiation**

| Stage of differentiation | Day | Media type | Compound | Final concentration |
|---|---|---|---|---|
| Mesodermal induction | 0 | CDM w/o insulin + BSA | Insulin (Sigma I9278) | 0.35ng/mL |
| | | | Fgf2 (Dr. Marko Hyvönen, Cambridge University) | 40ng/mL |
| | | | Chiron (Sigma CHIR99021) | 8µM |
| Paraxial mesodermal induction | 2 | CDM w/o insulin + BSA | Insulin | 7ng/mL |
| | | | Fgf2 | 4ng/mL |
| | | | Retinoic acid (Sigma R2625) | 1µM |
| Progenitor induction | 4 | CDM w/o insulin + BSA | Insulin | 7ng/mL |
| | | | Fgf2 | 4ng/mL |
| | | | Chiron | 3µM |
| | | | LDN 193189 (Sigma SML0559) | 100nM |
| | | | Ascorbic acid (Sigma A4403) | 10mg/ml |
| Adipose induction 1 | 6-8 | DMEM high glucose / HAM F12 (v/v) media<br>GlutaMAX (1/100e)<br>5% FCS<br>HEPES 15mM | T3 (Sigma T6397) | 1nM |
| | | | Dexamethasone (Sigma D4902) | 100nM |
| | | | IBMX (Sigma I7018) | 0.25mM |
| | | | Biotin (Sigma B4639) | 33µM |
| | | | Pantothenate (Santa Cruz SC278919) | 17µM |
| | | | Insulin | 500nM |
| | | | Rosiglitazone (Sigma R2408) | 5µM |
| | | | Ascorbic acid | 10mg/ml |
| Adipose induction 2 | 10 | DMEM high glucose / HAM F12 (v/v) media<br>GlutaMAX (1/100e)<br>5% FCS<br>HEPES 15mM | T3 | 1nM |
| | | | Dexamethasone | 100nM |
| | | | Biotin | 33µM |
| | | | Pantothenate | 17µM |
| | | | Insulin | 500nM |
| | | | Rosiglitasone | 5µM |
| Adipose maintenance | 12 – 25 | DMEM high glucose / HAM F12 (v/v) media<br>GlutaMAX (1/100e)<br>5% FCS<br>HEPES 15mM | T3 | 1nM |
| | | | Dexamethasone | 100nM |
| | | | Biotin | 33µM |
| | | | Pantothenate | 17µM |
| | | | Insulin | 500nM |
| | | | Rosiglitasone | 5µM |
| | | | Oleate (Sigma O1008) | 100µM |

**Table S3. qPCR primers list**

| Primer | Forward Sequence | Reverse Sequence |
|---|---|---|
| ADRβ3 | CTCGACGGGGCTTCTTGG | GAGGCCAGAGGTTTTCCACA |
| ADRP1 | GAGTCGTCTTCGGGACGCGC | TTGGCAACTGCAATTTGCGGC |
| CD36 | TGGAACAGAGGCTGACAACTT | TTGATTTTGATAGATATGGGATGC |
| C/EBP-α | GACATCAGCGCCTACATCG | GGCTGTGCTGGAACAGGT |
| C/EBP-β | CCAGCCCCCTCACTAATAGC | CCCTGCTCTGAGCTGTCG |
| C/EBP-δ | GGACATAGGAGCGCAAAGAA | GCTTCTCTCGCAGTTTAGTGG |
| DIO2 | CCTCCTCGATGCCTACAAAC | GCTGGCAAAGTCAAGAAGGT |
| EBF2 | AAGACCAACAACGGCACTCA | TTCGCAGCATCGACTACACA |
| GAPDH | AGCCACATCGCTCAGACAC | GCCCAATACGACCAAATCC |
| MYF5 | CTGCCAGTTCTCACCTTCTGA | AACTCGTCCCCAAATTCACCC |
| NANOG | ATGCCTCACACGGAGACTGT | CAGGGCTGTCCTGAATAAGC |
| OCT4 | GCTTCAAGAACATGTGTAAGCTG | AGGGTTTCCGTTTGCAT |
| PDGFRα | CCACCTGAGTGAGATTGTGG | TCTTCAGGAAGTCCAGGTGAA |
| PAX3 | ATTGGCAATGGCCTCTCA | AGGGGAGAGCGCGTAATC |
| PLIN1 | AGGGAAGAAGTTGAAGCTTGAGG | TTCTGGAAGCATTCGCAGGT |
| PPARα | GCACTGGAACTGGATGACAG | TTTAGAAGGCCAGGACGATCT |
| PPARγ | CGTGGCCGCAGATTTGAAAG | CACGGAGCTGATCCCAAAGT |
| PRDM16 | TGGCTGCTTCTGGACTCA | ATATTATTTACAACGTCACCGTCACT |
| SOX2 | GGGGGAATGGACCTTGTATAG | GCAAAGCTCCTACCGTACCA |
| TBOX | GCTGTGACAGGTACCCAACC | CATGCAGGTGAGTTGTCAGAA |
| UCP1 | CTCACCGCAGGGAAAGAA | GGTTGCCCAATGAATACTGC |
| ZIC1 | ATCCACAAAAGGACGCACAC | GTCACAGCCCTCAAACTCG |

**Table S4. Antibodies and dyes list**

| Antibody/Dye | Supplier | Identifier |
|---|---|---|
| AKT | CST | 9272S |
| C/EBPα | Santa Cruz | 14AA |
| COXII (MTCO2 12C4F12) | Invitrogen | A-6404 |
| DIO2 | Abcam | ab77779 |
| IRS1 | CST | 2382S |
| Ki-67 (8D5) | CST | 9449 |
| LipidTOX Deep Red neutral lipid stain | Thermofisher Scientific | H34477 |
| LipidTOX Green neutral lipid stain | Thermofisher Scientific | H34475 |
| LipidTOX red neutral lipid stain | Thermofisher Scientific | H34476 |
| Mitotracker™ Red CMXRos | Thermofisher Scientific | M7512 |
| Anti-mouse IgG, HRP-linked Antibody | CST | 7076S |
| MYF5 | Santa Cruz | sc302 |
| p-AKT | CST | 4051S |
| p-IRS1 | CST | 3070S |
| p-P70S6K | CST | 9206S |
| P70S6K | CST | 2708T |
| PAX3 | DSHB | AB528426 |
| PDGFRα (D13C6) XP | CST | 5241 |
| PGC1α | Abcam | ab54481 |
| PLIN1 | Progen | GP29 |
| PPARα (H98) | Santa Cruz | sc-9000 |
| PPARγ (E-8) | Santa Cruz | sc-7273 |
| PRDM16 | Abcam | ab106410 |
| Anti-rabbit IgG, HRP-linked Antibody | CST | 7074S |
| ß-actin | Abcam | ab16039 |
| TBOX | R&D | AF2085 |
| UCP1 | Abcam | ab155117 |
| UCP1 | Sigma | U6382 |
| ZIC1 | Abcam | 134951 |
| ADRP1 | Abcam | ab108323 |
